# A structural MRI marker predicts individual differences in impulsivity and classifies patients with behavioral-variant frontotemporal dementia from matched controls

**DOI:** 10.1101/2024.09.12.612706

**Authors:** Valérie Godefroy, Anais Durand, Marie-Christine Simon, Bernd Weber, Joseph Kable, Caryn Lerman, Fredrik Bergström, Maartje Luijten, Martine Groefsema, Guillaume Sescousse, Raffaella Migliaccio, Richard Levy, Bénédicte Batrancourt, Liane Schmidt, Hilke Plassmann, Leonie Koban

## Abstract

Impulsive decision-making is a symptom across many neuropsychiatric and neurological disorders. Yet, it is still unknown whether impulsivity can be predicted based on individual differences in brain structure. Here, we used machine-learning to develop a structural MRI signature of impulsivity, tested its validity in several independent samples, and assessed its diagnostic value in patients with behavioral variant frontotemporal dementia (bvFTD)—a neurodegenerative disease characterized by high impulsivity. The resulting whole-brain grey matter density pattern—the Structural Impulsivity Signature (SIS)—showed a good prediction-outcome correlation of *r*=0.35 (out-of-sample, *p*=0.0028) in the training sample of healthy adults (*N*=117) and significantly predicted individual differences in impulsivity in four other, independent studies (total *N*=626), including healthy and clinical participants with neuropsychiatric and neurological conditions. Further, the SIS separated bvFTD patients from controls with high accuracy (81% correct, *p*=0.002) and predicted impulsivity-related symptom severity among patients. The spatial distribution of weights in the SIS brain pattern highlights the key role of brain regions associated with affective processing in impulsivity. Together, these results provide evidence for a new structural neuromarker of individual differences in impulsivity that can be easily applied to new samples. Future studies can further assess its predictive value towards better prevention, diagnosis and treatment of mental disorders associated with impulsivity symptoms.

## Introduction

Impulsivity is the tendency to act in a rush and to seek immediate rewards without consideration of potentially negative consequences (Chamberlain and Sahakian, 2007). Trait impulsivity varies substantially within the general population, with high impulsivity being a hallmark of psychiatric and neurological conditions (Chamberlain et al., 2018). Despite the many negative consequences of high impulsivity for health and life in general (Goodwin et al., 2017; Rogers et al., 2021), its neurobiological correlates are still unclear, and it is unknown whether individual differences in impulsivity can be reliably predicted based on structural brain features (Bjork et al., 2009; Pehlivanova et al., 2018; MacNiven et al., 2020). Neurobiological measures of impulsivity could help to disentangle the heterogeneity of disorders related to maladaptive behavior and decision-making. Brain signatures of impulsivity could also constitute new predictive, diagnostic and prognostic biomarkers of conditions marked by impulsivity symptoms. They might in particular aid in the diagnosis, phenotyping and monitoring of conditions such as behavioral variant frontotemporal dementia (bvFTD)—a neurodegenerative disorder characterized by frontal and temporal brain atrophy, with high impulsivity and inappropriate behaviors as core symptoms (Rascovsky et al., 2011). Here, we aimed at developing a structural brain signature that could reliably predict individual differences in impulsivity in both healthy and clinical populations. Moreover, we tested this brain signature as a candidate diagnostic biomarker of bvFTD.

It has been shown that different types of measures load onto one latent construct of impulsivity (Chamberlain et al., 2018, 2019; Huang et al., 2024). These include cognitive tasks, personality trait questionnaires, and clinical measures of impulsivity-related symptoms. These findings suggest the existence of a coherent general dimension of impulsivity—characterized by an urgency to react—that can influence a range of behaviors across both healthy and clinical populations. At the same time, impulsivity is not a unitary construct; it can manifest through various mechanisms and be captured by different measures, such as delay discounting, which reflects the preference for smaller, immediate rewards over larger, delayed ones, or the tendency to pursue immediate gratification without regard for long-term consequences (Nombela et al., 2014).

Among measures of impulsivity, the behavioral measure of delay discounting may both reflect a personality trait and relate to clinical symptoms. Individual differences in delay discounting are relatively stable over time and show significant genetic heritability (Kirby, 2009; Anokhin et al., 2015; Koban et al., 2023). Moreover delay discounting is altered across many psychiatric (Amlung et al., 2019) and neurodegenerative (Godefroy et al., 2023) conditions often associated with high impulsivity. Recent studies have therefore started to investigate the neurobiological basis of individual differences in delay discounting, especially in healthy populations (Ballard and Knutson, 2009; Cooper et al., 2013; Lebreton et al., 2013; Li et al., 2013; Hare et al., 2014; van den Bos et al., 2014; Pehlivanova et al., 2018; Koban et al., 2023; Bergström et al., 2024). These studies, which have mostly used functional brain imaging with univariate approaches, have identified several networks which play a key role in delay discounting: the valuation and reward system (comprising the orbitofrontal cortex and ventromedial prefrontal cortex), the executive control system (including the dorsolateral prefrontal cortex), the memory and prospection system (including the hippocampus and medial temporal lobe).

It is very unlikely that complex psychological constructs such as impulsivity depend on only one or a few isolated brain regions. *Brain signatures* (or “neuromarkers”) go beyond the traditional univariate brain mapping approach by identifying multivariate *brain patterns* instead of examining brain regions independently (Genon et al., 2022) as potential correlates of impulsivity. Brain signatures are predictive models of mental events or of individual variables (such as impulsivity) that take into account distributed information across multiple brain systems (Kragel et al., 2018). Structural brain signatures are increasingly used in the field of translational neuroimaging, especially for applications in patients with neurodegenerative conditions (Woo et al., 2017). One of the greatest advantages of these predictive brain models is that they can be tested across studies, labs and populations to challenge their generalizability. We used this brain signature approach to identify a network of spatially distributed structural features predictive of differences in impulsivity in both healthy persons and persons with neuropsychiatric conditions. In addition, we aimed at testing the clinical value of this brain signature to detect a condition with core impulsivity symptoms due to brain structure damage.

Characterized by multiple impulsivity-related symptoms and significant structural modifications due to neurodegeneration, bvFTD is an ideal condition to demonstrate the diagnostic potential of a structural brain signature of impulsivity. BvFTD is the most common clinical variant of syndromes associated with predominant degeneration of the prefrontal and temporal regions as well as the basal ganglia. It is characterized by significant changes in personality and behavior including inhibition deficits (socially inappropriate and generally impulsive behaviors, compulsive stereotypic behaviors), as well as executive function deficits (Rascovsky et al., 2011). Most studies show alterations of delay discounting in bvFTD patients compared to controls (Bertoux et al., 2015; Chiong et al., 2016; Beagle et al., 2020; Manuel et al., 2020; Godefroy et al., 2023). Relatedly, brain regions known to be linked to impulsivity such as the orbitofrontal cortex (OFC), ventromedial prefrontal cortex (vmPFC) and ventral striatum (Levy and Glimcher, 2012; Rangel and Clithero, 2012; Bartra et al., 2013) are often affected in bvFTD (Chare et al., 2014; Karageorgiou and Miller, 2014).

Here, we first trained and cross-validated a structural MRI-based brain signature in a healthy adult population (Study 1, *N*=117) using LASSO-PCR (least absolute shrinkage and selection operator-principal component regression)—an established machine-learning algorithm (Tibshirani, 1996; Han et al., 2021)—to predict individual differences in impulsivity, as measured by delay discounting rates, based on participants’ grey matter maps (*N*=117). Brain markers of individual differences should ideally be tested in different and independent samples to establish their robustness and generalizability (Kragel et al., 2018). Thus, in Study 2 (*N*=166) and Study 3 (*N*=153), we tested the replicability of the brain signature in independent samples of healthy adults and with different validated measures of impulsivity (delay discounting and trait urgency in Study 2, trait impulsiveness in Study 3). In Study 4 (*N*=265), we tested the generalizability of brain-based predictions of impulsivity across healthy controls and participants with different neuropsychiatric conditions often associated with high impulsivity (schizophrenia, bipolar disorder and attention deficit hyperactivity disorder) (Chamorro et al., 2012). In Study 5, we tested the clinical value of the structural brain signature as a diagnostic biomarker in a population of patients with bvFTD, characterized by structural brain atrophy and core impulsivity symptoms (*N* = 42, including 24 bvFTD patients and 18 matched controls) (Godefroy et al., 2024). If a consistent pattern of grey matter density across the brain can reliably predict impulsivity, then the brain-predicted impulsivity should be higher in bvFTD patients than in controls and should be related to the level of clinically assessed impulsivity among patients. Additionally, as the neuroanatomical bases of impulsivity are still unclear, we analyzed the topographical distribution of the most important structural regions contributing to predict differences of impulsivity in healthy adults (in Study 1) and in bvFTD (in Study 5).

## Methods and Materials

We used the data collected in a total of five studies to develop and test the brain signature of impulsivity (see Fig. 1.A). This allowed us to: 1) develop a structural brain signature of trait impulsivity—which we refer to as the *Structural Impulsivity Signature (SIS)*—in healthy adults (Study 1); 2) test its validity in independent samples of healthy and clinical populations (neuropsychiatric conditions often associated with high impulsivity) using various measures of impulsivity trait (Studies 2, 3, 4), and 3) test its validity and clinical value for diagnosis in bvFTD, a neurodegenerative condition in which brain structure lesions contribute to impulsivity symptoms (Study 5). Results from all tested datasets are presented in this manuscript.

**Figure 1.**
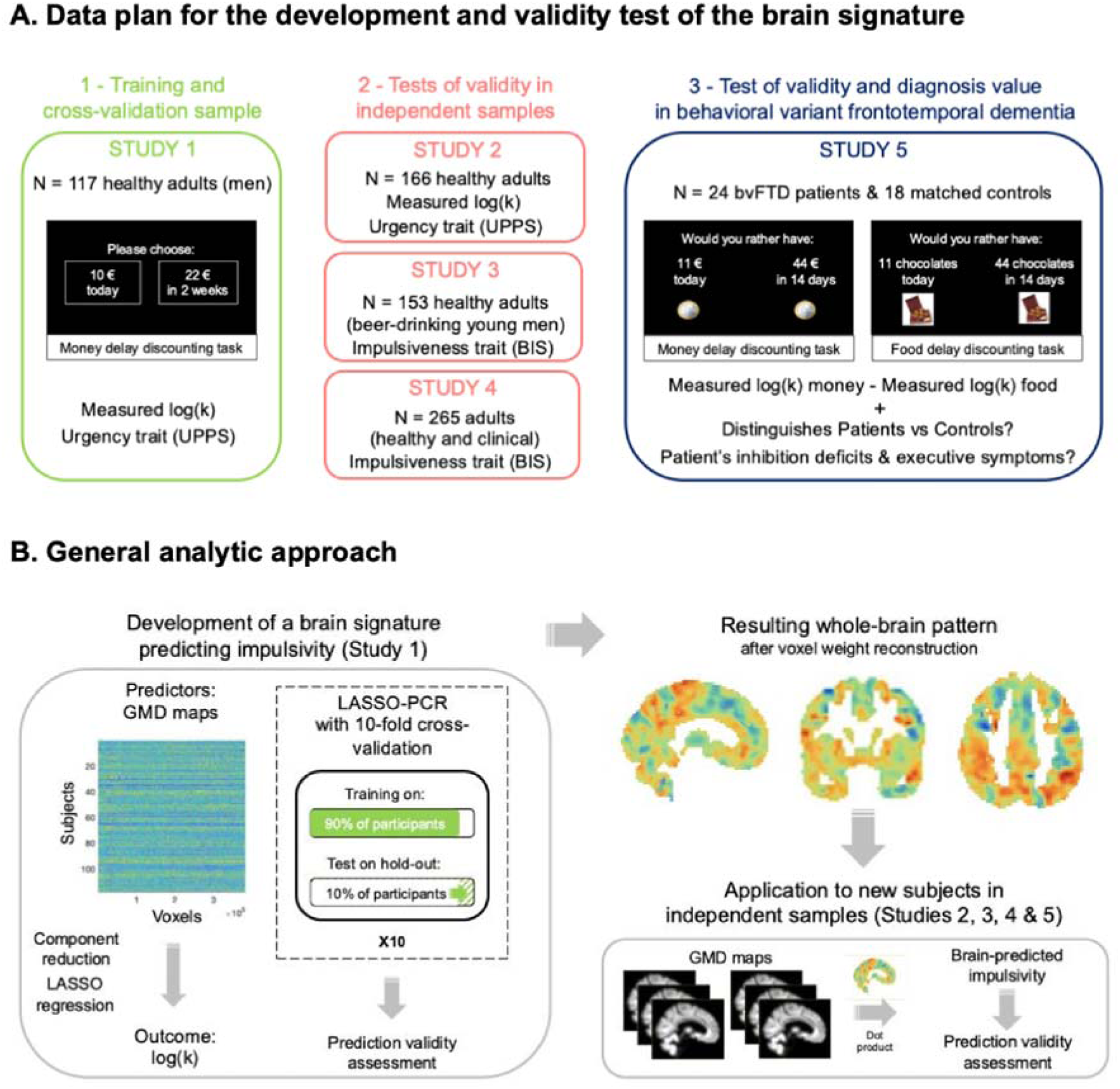
Methodological approach for the development and validation of a structural brain signature of impulsivity. **(A)** Several tests were performed in each of the five studies to assess the validity of the structural signature trained in Study 1. In Study 1 (the training and cross-validation sample), permutation tests on different metrics (*MSE, RMSE, MAE*) and on the correlation between the Structural Impulsivity Signature (SIS) response and measured delay discounting (*log(k)*) were used to investigate the predictive accuracy of the developed brain pattern; the validity of SIS responses was also assessed through testing their correlation with out-of-sample *log(k)* measured several weeks later and with a self-report measure of urgency trait from the Urgency, lack of Premeditation, lack of Perseverance, and Sensation seeking (UPPS) model of impulsive behaviors. Studies 2, 3, 4 served as independent test samples to further validate and generalize the structural signature developed in Study 1; Study 5 was an additional independent sample to test the clinical value of the structural signature for diagnosis. In Study 2, we tested whether SIS responses correlated with *log(k)*’s and with self-reported urgency trait (from the UPPS model of impulsive behaviors) in healthy adults. In Study 3 and Study 4, we tested whether SIS responses correlated with impulsiveness trait (from Barratt Impulsiveness Scale) in young male adults (Study 3) and in a mixed sample of healthy controls and participants with different psychiatric conditions (schizophrenia, bipolar disorder and attention deficit hyperactivity disorder - Study 4). In Study 5, which involved patients with behavioral variant frontotemporal dementia (bvFTD) matched with healthy controls, we tested: 1) correlations between SIS responses and observed delay discounting for two types of stimuli (money and food) across patients and controls; 2) the ability of SIS responses to distinguish patients from controls; 3) correlations between measures of impulsivity symptoms (inhibition and executive deficits) and SIS responses among patients. **(B)** Grey matter density (GMD) maps from healthy participants of Study 1 were used for the prediction of delay discounting (*log(k)*) by LASSO-PCR with 10-fold cross-validation. In each fold, the classifier was trained on 90% of the data and tested on the remaining 10% hold-out data to evaluate its predictive accuracy. The resulting brain pattern was applied to the grey matter density maps of each study’s participants to evaluate the validity of its predictions in different types of population and with different measures of impulsivity (Studies 2 to 5).

### Participants

The research reported here complies with all relevant ethical regulations. The study protocols were approved by the institutional review board of Bonn University’s Medical School (Study 1), by the University of Pennsylvania Institutional Review Board (Study 2), by the regional ethics committee CMO-Arnhem-Nijmegen (Study 3), by the University of California Los Angeles Institutional Review Board (Study 4), and by the French Ethics Committee “Comité de Protection des Personnes Sud Méditerranée I” (Study 5).

### Study 1

In Study 1, participants were recruited in the context of a seven-week dietary intervention study (https://osf.io/rj8sw/?view_only=af9cba7f84064e61b29757f768a8d3bf) at the University of Bonn in Germany. In this study, only male participants were recruited to exclude potential menstrual cycle effects, with the following inclusion criteria: age between 20 and 60 years, right-handedness, non-smoker, no excessive drug or alcohol use in the past year, no psychiatric or neurological disease, body mass index (*BMI*) between 20 and 34, no other chronic illness or medication, following a typical Western diet without dietary restrictions, and no MRI exclusion criteria (e.g., large tattoos, metal in the body). For the present purpose, we used only the behavioral and structural MRI data collected before the dietary intervention. *N*=117 participants were tested for the baseline session of Study 1. However, four participants were excluded from the present analyses due to being outliers on grey matter density maps (three participants) and due to very inconsistent choices at the intertemporal choice task (one participant). Thus, the data of a total of 113 participants was used for the analyses.

### Study 2

In Study 2, participants were recruited in the context of a ten-week cognitive training study (registered at clinicaltrials.gov as Clinical trial reg. no. NCT01252966) at the University of Pennsylvania, USA. Individuals between 18 and 35 years of age who reported home computer and internet access were recruited. Exclusion criteria were: an IQ score of <90 on Shipley Institute of Living Scale, self-reported history of neurological, psychiatric, or addictive disorders (excluding nicotine), positive breath alcohol reading (>0.01), color blindness, left-handedness, and claustrophobia. Here, we focused on behavioral and structural MRI data collected during the baseline session before the cognitive training. In Study 2, *N*=166 participants (mean age=24.5, 59% male, 41% female) were included in the baseline session and all were included in our data analyses.

### Study 3

In the context of a project on alcohol use in young adults(Groefsema et al., 2020b, 2020a) at Radboud University, participants were recruited using flyers and online advertisement. Inclusion criteria were as follows: (a) age 18–25, (b) drinking beer and (c) being male. Exclusion criteria were MRI contraindications and a history of brain injury. The final sample used for our analyses consisted of 153 young adults (mean age = 22.8 years). The recruited participants showed different levels of alcohol use based on two self-report measures, as well as the DSM-IV (Diagnostic and Statistical Manual of Mental Disorders-IV) criteria for alcohol dependence.

### Study 4

In the context of a large project on brain function and anatomy in common neuropsychiatric syndromes(Gorgolewski et al., 2017) led by the Consortium for Neuropsychiatric Phenomics, participants were recruited by community advertisement and through outreach to local clinics and online portals. Each subject had completed at least 8 years of formal education and had either English or Spanish as primary language. The consortium excluded patients with diagnoses in at least 2 different patient groups. Moreover, the following exclusion criteria were used: left-handedness, pregnancy, history of head injury with loss of consciousness or other contraindications to scanning. The consortium published a dataset(Poldrack et al., 2016) with neuroimaging as well as phenotypic information for 272 participants (mean age=33.3, 57.7% male). The final sample used for our analyses consisted of a total of 265 participants with available T1 weighted scans, including healthy adults (*N*=125), patients with schizophrenia (*N*=50), patients with bipolar disorders (*N*=49) and patients with attention deficit hyperactivity disorder (*N*=41).

### Study 5

For Study 5, participants were recruited in the context of a clinical study at the Paris Brain Institute, France (clinicaltrials.gov: NCT03272230). This study was designed to investigate the behavioral correlates and neural bases of neuropsychiatric symptoms associated with behavioral variant frontotemporal dementia (bvFTD). BvFTD patients were recruited in two tertiary referral centers, at the Pitié-Salpe-trière Hospital and the Lariboisière Fernand-Widal Hospital, in Paris. Patients were diagnosed according to the International Consensus Diagnostic Criteria(Rascovsky et al., 2011). To be included, bvFTD patients had to present a Mini-Mental State Evaluation (MMSE) score of at least 20. Healthy controls (HC) were recruited by an online announcement. Inclusion criteria included a MMSE score of at least 27 and matching the demographic characteristics of the bvFTD group. In total, we used the data of 24 bvFTD patients (mean age=66.6, 66.6% male, mean disease duration = 3.8 years) and 18 controls matched to patients for age and sex (mean age=62.6, 44.4% male) in this clinical study (see Supplementary table 1).

As detailed in Figure 1A, we tested the generalizability of SIS responses not only in substantially different populations (e.g., from different labs, regions, ages, healthy and clinical) but also with different types of measures of impulsivity: delay discounting measures from intertemporal choice tasks (Studies 1, 2 and 5), trait measures by two different questionnaires (Studies 1, 2, 3 and 4) and clinical measures of symptoms (Study 5).

### Intertemporal choice tasks

#### Study 1

During the intertemporal choice (ITC) task performed in an MRI scanner, participants in Study 1 were presented with 108 trials offering a choice between a smaller sooner (SS) reward option and a larger later (LL) reward option(Koban et al., 2023). Participants were informed that one of their choices could be paid out at the end of the experiment, which made their choices non-hypothetical and incentive-compatible. Participants used their left or right index finger to press the response key corresponding to their choice (left index for left option or right index for right option). The option chosen by the participant was then highlighted by a yellow frame (see Koban et al., 2023(Koban et al., 2023) for further details on the trial structure).

#### Study 2

During the ITC performed in an MRI scanner, participants had to make 120 choices between the same smaller immediate reward ($20 today) and a varying larger reward available after a longer delay (e.g., $40 in a month)(Kable et al., 2017). Participants were informed that one of their choices could be paid out at the end of the experiment, which made their choices non-hypothetical and incentive-compatible. Each trial started with the presentation of the amount and delay of the larger later option. Once subjects had made their choice, a checkmark on the screen indicated if the larger later option was chosen and a “X” indicated that the immediate option was chosen (see Koban et al., 2023(Koban et al., 2023) for further details on the trial structure).

#### Study 5

In Study 5, participants performed two ITC tasks on a computer screen, one using monetary rewards (from 8 to 35 euros) and one using food rewards (from 8 to 35 chocolates) in randomized order(Godefroy et al., 2024). Each of these tasks included 32 choices between SS and LL options. Like in Study 1 and 2, participants’ choices were non-hypothetical and incentive-compatible. For each trial, participants could indicate their choice by pressing either a blue key on the keyboard with their right-hand index to select the option on the left or a yellow key with their right-hand middle finger to select the option on the right. Once the choice had been made, a message on the screen indicated which option had been chosen (see Godefroy et al. (Godefroy et al., 2024) for further details on the trial structure).

### Other measures of impulsivity traits and impulsivity symptoms

#### Study 1

In Study 1, along with choice data collected from the ITC task, we used questionnaire data from the Impulsive Behavior Short Scale–8 (I-8), which measures impulsivity traits according to the Urgency, lack of Premeditation, lack of Perseverance, and Sensation seeking (UPPS) model with four subscales comprising two items each(Groskurth et al., 2022). We predicted that the trait of *urgency— defined as the tendency to act rashly in an emotional context* (e.g., “I sometimes do things to cheer myself up that I later regret”)*—* would be closest to SIS responses, supposed to measure impulsivity as the urgency to react, the tendency to choose the most prepotent behavioral answers to any situation without considering potentially negative consequences.

#### Study 2

In Study 2, we used data from the UPPS-P Impulsive Behavior Scale, which measures trait impulsivity according to the UPPS model(Whiteside and Lynam, 2001) like in Study 1. We also predicted that urgency trait would be the most closely related to SIS responses. We used the average of the subscales of positive urgency (rash actions taken in response to positive emotional states) and negative urgency (rash actions taken in response to negative emotional states) to test this hypothesis.

#### Study 3

Study 3 used yet a different measure of impulsivity based on the Barratt Impulsiveness Scale (BIS), a 30-item self-report instrument designed to assess the personality construct of impulsiveness defined “as a predisposition toward rapid, unplanned reactions to internal or external stimuli without regard to the negative consequences of these reactions to the impulsive individuals or to others”(Patton et al., 1995; Stanford et al., 2009). We predicted that trait impulsivity measured by BIS would be related to the SIS responses across the whole sample, including participants with different levels of alcohol dependence. We used the sum of the scores on the three BIS subscales (attentional impulsiveness, motor impulsiveness, non-planning impulsiveness) as the measure of impulsivity to test this hypothesis.

#### Study 4

In Study 4, we also used data from the Barratt Impulsiveness Scale (BIS)(Patton et al., 1995; Stanford et al., 2009). We predicted that the trait impulsivity measured by BIS (sum of the scores on the three subscales) would be related to SIS responses across the whole sample, including both healthy adults and patients with neuropsychiatric syndromes.

#### Study 5

In this study, we used clinical measures of core impulsivity symptoms of bvFTD, in particular: inhibition deficit and dysexecutive syndrome. Another recent investigation of the same sample found that these two bvFTD symptoms are related to higher discounting rates(Godefroy et al., 2024). As an objective measure of inhibition deficit, we used the Hayling Sentence Completion Test (HSCT)(Burgess and Shallice, 1997). In the HSCT, participants are asked to complete 15 sentences using the appropriate word, as fast as possible (automatic condition, part A), and 15 sentences using a completely unrelated word (inhibition condition, part B). We used the Hayling error score (number of errors in part B) as a measure of the difficulty to inhibit a prepotent response, as in Flanagan et al.(Flanagan et al., 2016). As a measure of executive functions, we used the Frontal Assessment Battery (FAB)(Dubois et al., 2000) (lower scores indicating worse executive functions).

### MRI data acquisition and preprocessing

#### Study 1

Brain imaging data for Study 1 were acquired using a Siemens Trio 3T scanner. Structural images were acquired using a T1 weighted MPRAGE sequence with the following parameters: TR 1660 ms; TE 2.54 ms; FoV 256 mm; 208 slices; slice thickness 0.80 mm; TI 850 ms; flip angle 9°; voxel size 0.8 mm isomorphic; total acquisition time 6:32 min. T1 images were preprocessed for Voxel Based Morphometry (VBM) analyses with SPM 12 using parameters described in the Supplementary Material. To optimize the sample homogeneity among the obtained grey matter images, three outliers (based on Mahalanobis distance of individual grey matter density maps with Bonferroni correction(Ghorbani, 2019)) were detected and excluded from further analyses.

#### Study 2

Brain imaging data for Study 2 were acquired using a Siemens Trio 3T scanner (with a 32-channel head coil). Structural images were acquired using a T1 weighted MPRAGE sequence with the following parameters: TR 1630 ms; TE 3.11 ms; FOV 192x256; 160 slices; slice thickness 1 mm; TI 1100 ms; flip angle 15°; voxel size 0.9375 × 0.9375 × 1.000 mm; total acquisition time 4:35 min. We used existing data preprocessed by Kable and colleagues(Kable et al., 2017) (see Supplementary Material).

#### Study 3

Brain imaging for Study 3 was conducted on a PRISMA(Fit) 3T Siemens scanner, using a 32-channel head coil. A T1 weighted MPRAGE sequence was acquired in each participant with the following parameters: TR 300 ms; TE 3.03 ms; FOV 256 mm; 192 slices; voxel size 1.0 × 1.0 × 1.0 mm. T1 images were preprocessed for VBM analyses using SPM 12 (see Supplementary Material).

#### Study 4

Brain imaging data for Study 4 was obtained from the OpenfMRI database (accession number ds000030). Images were acquired using a Siemens Trio 3T scanner. T1-weighted high-resolution anatomical scans (MPRAGE) were collected with the following parameters: TR 1.9s; TE 2.26ms; FOV 250mm; 176 slices; slice thickness 1mm. T1 images were preprocessed for VBM analyses using SPM 12 (see Supplementary Material).

#### Study 5

Brain imaging data for Study 5 were acquired using a Siemens Prisma whole-body 3T scanner (with a 12-channel head coil). Structural images were acquired using a T1 weighted MPRAGE sequence with the following parameters: TR 2400 ms; TE 2.17 ms; FOV 224 mm; 256 slices; slice thickness 0.70 mm; TI 1000 ms; flip angle 8°; voxel size 0.7 mm isomorphic; total acquisition time 7:38 min. T1 images were preprocessed for VBM analyses using SPM 12 (see Supplementary Material).

### Data analyses

The global analytic approach is summarized in Figure 1B. We used a study with a sample size superior to a hundred participants (Study 1) for the development of the Structural Impulsivity Signature (SIS). Most recent studies using the brain signature framework suggest that sufficient in-sample power and out-of-sample replication probability can be achieved for a variety of phenotypes for this sample size(Spisak et al., 2022). Some behavioral data were missing for a few participants in Study 5 (*N*=4 missing data for *log(k)’s* computed for monetary rewards and food rewards; *N*=1 missing data for the Hayling error score); these participants were therefore excluded of the correlation tests involving each these specific variables. All analyses were performed using R Studio (1.2.1335) and Matlab (R2017b).

### Computation of discount rates

In all three studies, the individual discounting rate (k) was estimated by fitting logistic regressions to the individual choice data, with the assumption that the subjective value (*SV*) of the choice options followed hyperbolic discounting, as follows:

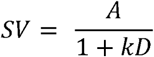

where *A* is the amount of the option, *D* is the delay until the receipt of the reward (for immediate choice, *D* = 0), and *k* is a discounting rate parameter that varies across subjects. Higher values of *k* indicate greater discounting and thus higher preference for sooner rewards. In Study 1, we used logistic regressions (as described in Wileyto et al., 2004) to estimate the individual parameter *k* from the participant’s answers in the ICT task at baseline and we used the *log(k)* values as the parameter to be predicted. Individual *k*’s were log-transformed in all studies to obtain non-skewed distributions of discounting parameters. In Study 2, we also used the *log(k)* values at baseline (see(Kable et al., 2017)). In Study 5, we used the *log(k)* values calculated in bvFTD patients matched with controls for both monetary and food rewards (see(Godefroy et al., 2024)).

### LASSO-PCR, training and cross-validation of the Structural Impulsivity Signature in Study 1

We used a standard machine learning algorithm, LASSO-PCR (least absolute shrinkage and selection operator-principal component regression) (Tibshirani, 1996), to train a classifier to predict *log(k)* from the individual whole brain grey matter density (GMD) maps. LASSO-PCR uses principal components analysis (PCA) to reduce the dimensionality of the data and LASSO regression to predict the outcome (*log(k)*) from the component scores. The components identified by the PCA correspond to groups of brain regions that covary with each other in terms of grey matter density. The LASSO algorithm fits a regularized regression model predicting *log(k)* from the identified components. This algorithm iteratively shrinks the regression weights towards zero, thus selecting a subset of predictors and reducing the contribution of unstable components. LASSO-PCR is suited to make predictions from thousands of voxels across the whole-brain, in particular because it reduces multicollinearity between voxels(Wager et al., 2011, 2013). It also allows reconstructing voxel weights across the brain (from voxel loadings on PCA components and LASSO regression coefficients of components), yielding predictive brain maps that are easier to interpret than component weights. To assess the accuracy of this predictive modeling from GMD maps, we used a 10-fold cross-validation process. The brain classifier was trained on 90% of the data and tested on the remaining 10% with 10 iterations, so that each participant was used for training the model in nine folds and for testing the accuracy of its prediction in the remaining fold. Ten-fold cross-validation is within the range of typically recommended folds (between 5 and 10) and allowed for a large training sample size at each iteration (Scheinost et al., 2019; Poldrack et al., 2020). Default regularization parameters were used for all machine-learning analyses to avoid overfitting of the model to the data. We used four metrics to assess the accuracy of the model predictions: the mean squared error (*MSE*) of prediction, the root mean squared error (*RMSE*), the mean absolute error (*MAE*), and the correlation between the model predictions (from the 10 hold-out test samples) and observed *log(k)*’s (prediction-outcome correlation).

### Validity of SIS responses in Study 1

To test the reliability of the predictions, we used permutation tests assessing the statistical significance of the accuracy metrics (*MSE*, *RMSE*, *MAE* and prediction-outcome correlation). More precisely, 5000 iterations of randomly permuting the *log(k)* values were used to generate null distributions of these four metrics and thus to assess the probability of: (*MSE* < actual *MSE*), (*RMSE* < actual *RMSE*), (Mean abs. error < actual Mean abs. error) and of (prediction-outcome correlation > actual prediction-outcome correlation) under the null hypothesis. To further confirm the validity of out-of-sample predictions of trait impulsivity, we performed correlation tests between the SIS responses and: (1) calculated log(k) values for the ITC task performed seven weeks later at the end of the dietary intervention (one-tailed test for a clear directional hypothesis of correlation); (2) the urgency trait subscale of the Impulsive Behavior Short Scale–8 (I-8).

### Validity of SIS responses in independent samples of healthy and clinical participants in Studies 2-4

To assess the predictions of the brain classifier developed in Study 1 in participants of Studies 2 to 4, we calculated the dot product between the predictive weight map (SIS) and the grey matter density map of each participant of Study 2, 3 and 4. The dot product (computed as a linear combination of the participant’s voxel grey matter density multiplied by voxel weight across the brain), plus the classifier’s intercept, provides a pattern response or SIS response and thereby a predicted value of trait impulsivity for each participant. This allowed us to test the correlations between the SIS responses and: (1) the *log(k)* values computed in the sample as well as the urgency trait (measured by the UPPS-P Impulsive Behavior Scale) in Study 2; (2) the impulsiveness trait (measured by the BIS scale) across the whole sample in Study 3 and in Study 4.

### Validity and diagnostic value of SIS responses in patients with bvFTD in Study 5

To assess the predictions of the brain classifier developed in Study 1 in participants of Study 5, we calculated again the dot product as a measure of SIS response and thereby a predicted value of impulsivity for each participant of Study 5. This allowed us to test: (1) the correlation between the SIS responses and the *log(k)* values (computed for both monetary and food rewards) across the whole sample (one-tailed tests for clear directional hypotheses of correlation); (2) whether the SIS responses could accurately discriminate between bvFTD patients and controls, using a single-interval test (thresholded for optimal overall accuracy); (3) whether the SIS responses were related to the severity of inhibition deficit (measured by Hayling error score) and of dysexecutive syndrome (i.e., lower FAB total score) among bvFTD patients.

### Spatial distribution of regions contributing to predict differences of impulsivity

We used a bootstrapping analysis to detect the brain regions that were the most robust contributors to predict impulsivity (measured by *log(k)* in Study 1). Sampling with replacement from the initial sample of Study 1 participants generated 5,000 samples. The LASSO-PCR algorithm yielded a predictive brain pattern (voxel weights across the brain) from the data (paired GMD map – *log(k)* outcome) in each of these 5,000 samples. For each voxel weight in the whole-brain pattern, the probability of being different from 0 (either above or below 0) could be estimated across the 5,000 samples. Thus, two-tailed, uncorrected *p*-values were calculated for each voxel across the whole brain and false discovery rate (FDR) correction was used to correct for multiple comparisons. Bootstrapped weights were thresholded at *q*=0.05 FDR-corrected across the whole weight map, as well as at *p*=0.05 uncorrected for display. To identify the main brain regions which contributed to differentiate bvFTD patients from controls on SIS response, we contrasted bvFTD patients versus controls in terms of voxel-wise pattern expression of the predictive map of impulsivity (see further details in result section).

### Data availability

Deidentified data of Studies 1, 2 and 5 are available at https://figshare.com/s/a090ef2531a7d7b9a2e4. Data from Study 3 are available upon request to the first author. Data from Study 4 are available at https://openfmri.org/dataset/ds000030/

## Results

### Development and cross-validation of the Structural Impulsivity Signature in healthy adults

In Study 1, on average, participants had a fitted *log(k)* parameter of -5.94 (median *log(k)*=-5.49, corresponding to *k*=0.0041). Discounting rates were characterized by substantial individual differences (*SD*=2.00), with *log(k)* ranging from -11.92 to -2.16. *Log(k)* showed a trend for a positive correlation with the urgency trait (*R*=0.17, *p*=0.06, 95%-*CI*= [-0.009, 0.35]).

The 10-fold cross-validation procedure revealed a significant accuracy of the Structural Impulsivity Signature (SIS) responses (see Figure 2A and 2B and Supplementary Figure 1): the predictions had a mean squared error of 3.45 (permutation test: *p*=0.0026), a root mean squared error of 1.86 (permutation test: *p*=0.0026), a mean absolute error for predicted *log(k)* of 1.46 (permutation test: *p*=0.0022), and a cross-validated prediction-outcome correlation of *R*=0.35 (permutation test: *p*=0.0028) (Figure 2C). Further, supporting the reliability and conceptual validity of the SIS responses, we found that they significantly correlated with (out-of-sample) *log(k)*’s computed from the same ITC task performed seven weeks later by the same participants (*R*=0.34, *p*<0.001, 95%-*CI*= [0.18, 1]). This suggests that a relatively stable part of the between-person variability in delay discounting was explained by individual differences in brain structure. Moreover, higher SIS responses were associated with higher self-reported urgency (*R*=0.20, *p*=0.037, 95%-*CI*= [0.01, 0.37]) (Figure 2D). For completeness, we checked that other subscales of the UPPS did not correlate with SIS responses. Indeed, SIS responses did not correlate with lack of planning (*R*= -0.04; *p*= 0.64), lack of perseverance (*R*= -0.01; *p*= 0.90) or sensation seeking (*R*= -0.01; *p*= 0.89).

**Figure 2.**
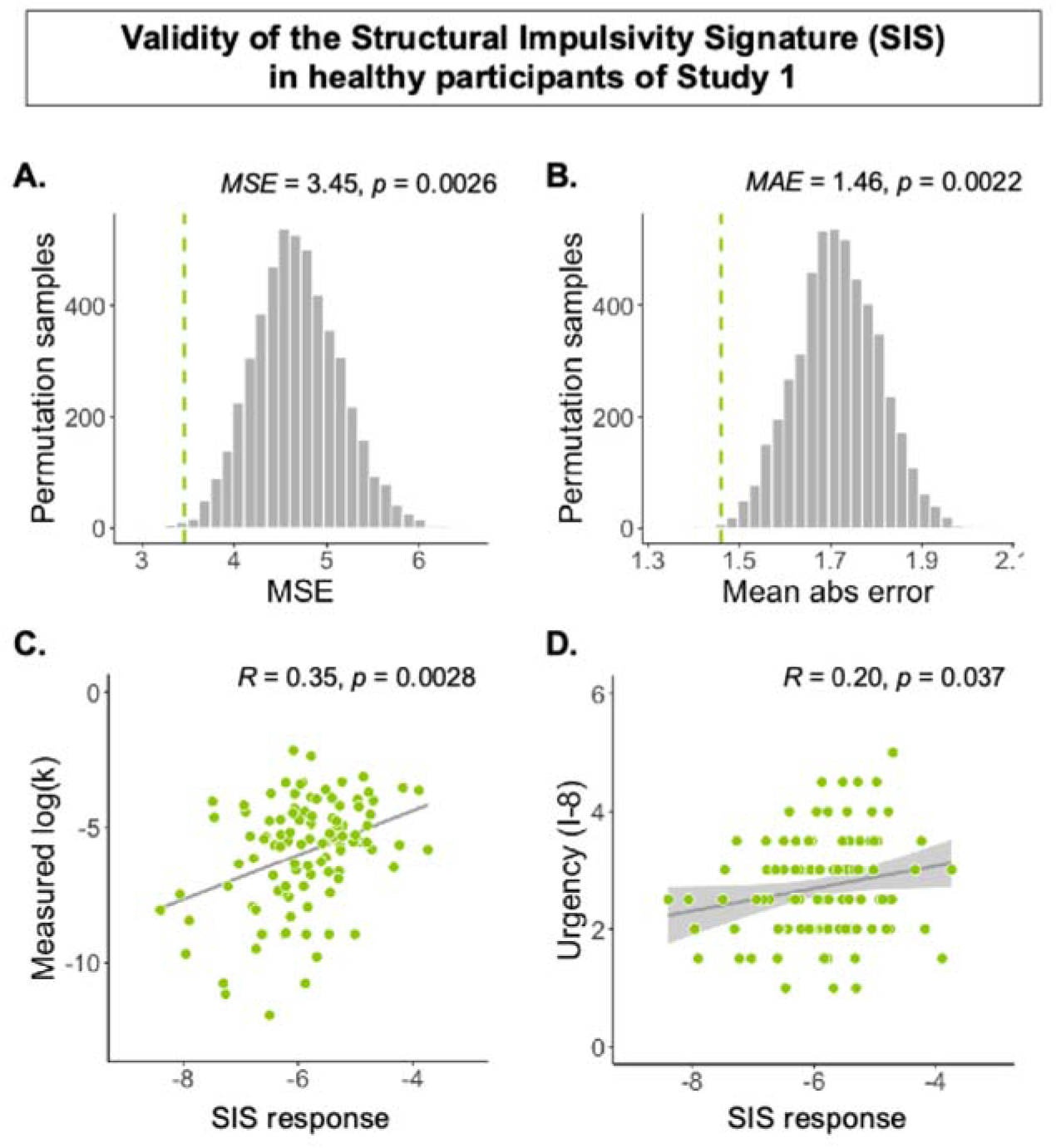
Predictive validity of the structural brain pattern in Study 1. **(A)** Mean squared error (*MSE*) of prediction and significance obtained by permutation test (5,000 samples – *N*=113 males). **(B)** Mean absolute error (*MAE*) of prediction and significance obtained by permutation test (5,000 samples – *N*=113 males). **(C)** Correlation between SIS responses and measured *log(k)*, with significance of prediction-outcome correlation obtained by permutation test (5,000 samples – *N*=113 males). **(D)** Test of the correlation between SIS responses and self-reported urgency (subscale of I-8 Impulsive Behavior Short Scale) (*R*=0.20, *p*=0.037, 95%-*CI*= [0.01, 0.37]).

Like the measures of log(k) in this sample(Koban et al., 2023), SIS responses did not significantly correlate with age (*R*=-0.11, *p*=0.24, 95%-*CI*= [-0.29, 0.07]), education (*R*=-0.15, *p*=0.10, 95%-*CI*= [-0.33, 0.03]), income (*R*=-0.12, *p*=0.21, 95%-*CI*= [-0.30, 0.07]), BMI (*R*= -0.04, *p*=0.66, 95%-*CI*= [-0.22, 0.14]), and percentage of body fat (*R*= -0.13, *p*=0.18, 95%-*CI*= [-0.31, 0.06]) (see more details in Supplementary Figure 2).

### Validity of the SIS in independent samples of healthy and clinical populations

Study 2 tests the predictions of the Structural Impulsivity Signature (SIS) in a second MRI dataset of healthy participants, that has used a different protocol, scanner, different preprocessing pipeline, in a socio-demographically different participant population. While we did not find a significant link between SIS response and observed *log(k)* in Study 2 (*R*=0.06, *p*=0.21, 95%-*CI*= [-0.07, 1]), SIS response was positively associated with urgency trait (*R*=0.15, *p*=0.047, 95%-*CI*= [0.002, 0.30], see Figure 3A), as in Study 1. Moreover, like in Study 1, SIS responses did not correlate with the other subscales of the UPPS. Of note, in Study 2, individual differences in the discounting parameter were less variable (*SD*=0.98) as compared to Study 1, with *log(k)* ranging from -7.08 to -2.12. Additionally, observed *log(k)* had a trend for a negative correlation with urgency (*R*=-0.14, *p*=0.06, 95%-*CI*= [-0.29, 0.008]). Therefore, in Study 2, the discounting rate does not seem to be related to individual differences in impulsivity trait.

**Figure 3.**
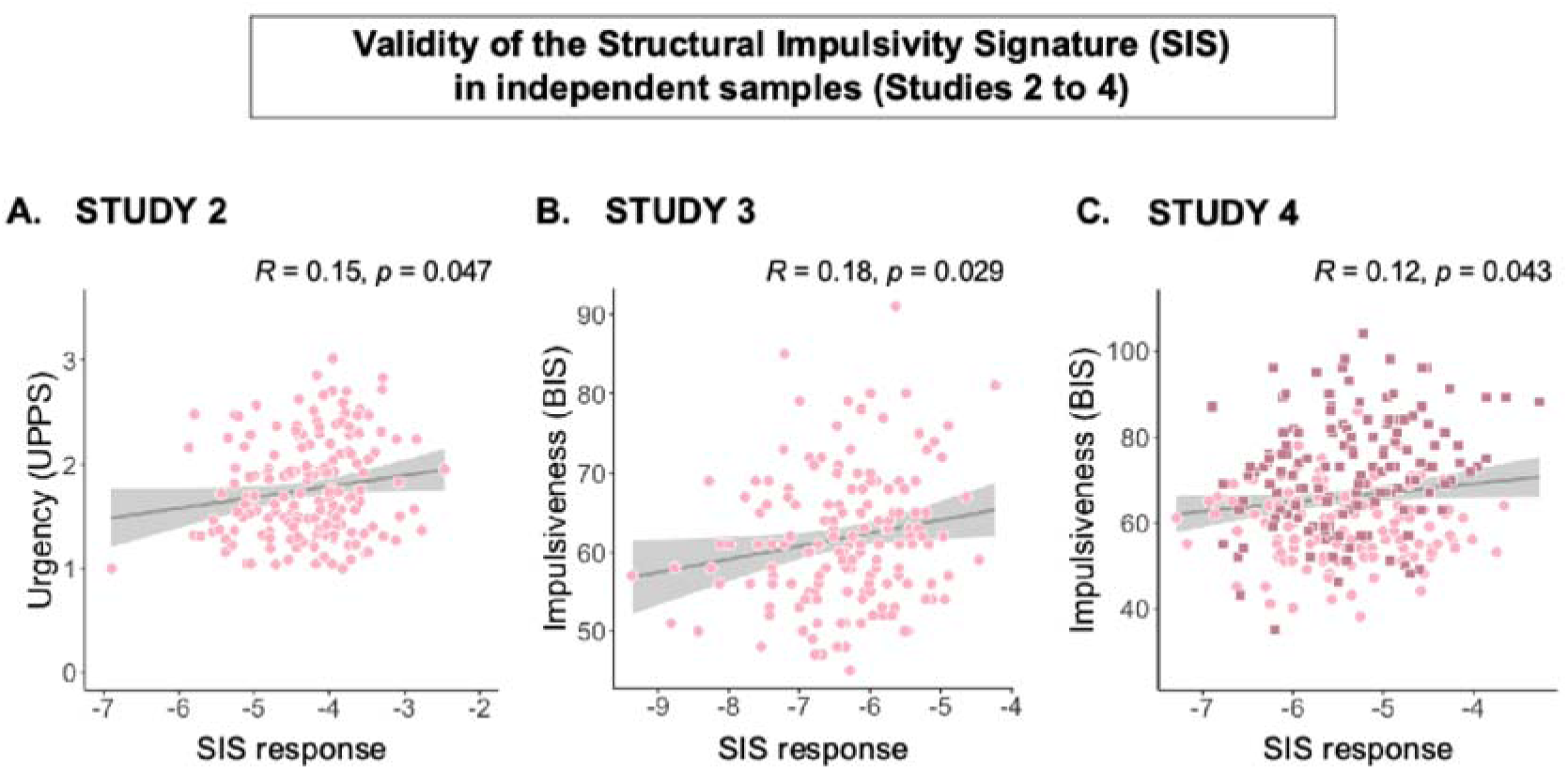
Predictive validity of the structural brain pattern in Study 2, Study 3 and Study 4. **(A)** Test of the correlation between SIS responses and self-reported urgency (mean of positive and negative urgency subscales of UPPS-P Impulsive Behavior Scale) in Study 2 (*R*=0.15, *p*=0.047, 95%-*CI*= [0.002, 0.30]). (**B)** Correlation between SIS responses and self-reported impulsiveness (total of Barratt Impulsiveness Scale) in Study 3 (*R*=0.18, *p*=0.029, 95%-*CI*= [0.02, 0.33]). **(C)** Correlation between SIS responses and self-reported impulsiveness (total of Barratt Impulsiveness Scale) in Study 4 in healthy participants (light pink circles) and patients with different psychiatric conditions (dark pink squares) (*R*=0.12, *p*=0.043, 95%-*CI*= [0.004, 0.24]).

Study 3 tests the predictions of the SIS in a population of young male adults with different levels of alcohol use. Study 4 tests the predictions of the SIS in a population including both healthy adults and patients with neuropsychiatric conditions (schizophrenia, bipolar disorder, and ADHD) in which high impulsivity is a common feature (Chamorro et al., 2012). SIS response was positively associated with impulsiveness trait across the whole sample in Study 3 (*R*=0.18, *p*=0.029, 95%-*CI*= [0.02, 0.33], see Figure 3B), as well as in Study 4 (*R*=0.12, *p*=0.043, 95%-*CI*= [0.004, 0.24], see Figure 3C). Moreover, in Study 4, SIS response significantly correlated with impulsiveness trait among patients (*R*=0.23, *p*=0.006, 95%-*CI*= [0.07, 0.38], see Figure 3D), which supports the idea that the SIS could have applications for precision medicine, for the characterization of impulsivity phenotypes in patients with neuropsychiatric conditions.

### Validity and diagnostic value of the SIS in a sample of bvFTD patients and matched controls

Study 5 allowed us to investigate the accuracy of SIS responses and the clinical relevance of the SIS (1) for classifying patients with bvFTD differently from matched control participants and (2) for predicting the core symptoms of disinhibition and executive deficits among patients with bvFTD (Rascovsky et al., 2011). In line with the core symptoms of this disorder, bvFTD patients presented significantly higher delay discounting (i.e. more impatient or impulsive choices) compared to controls, for both money rewards and food rewards(Godefroy et al., 2024). They also showed higher inhibition deficit and lower executive performances compared to controls (see Supplementary table 1).

To confirm the predictive validity of our classifier in Study 5, we showed that the SIS responses were positively correlated with *log(k)* values, for both monetary rewards (*R*=0.30, *p*=0.03, 95%-*CI*= [0.03, 1], mean absolute error of 2.08) and for food rewards (*R*=0.40, *p*=0.006, 95%-*CI*= [0.15, 1], mean absolute error of 2.65) (see Figure 4.A and 4.B).

**Figure 4.**
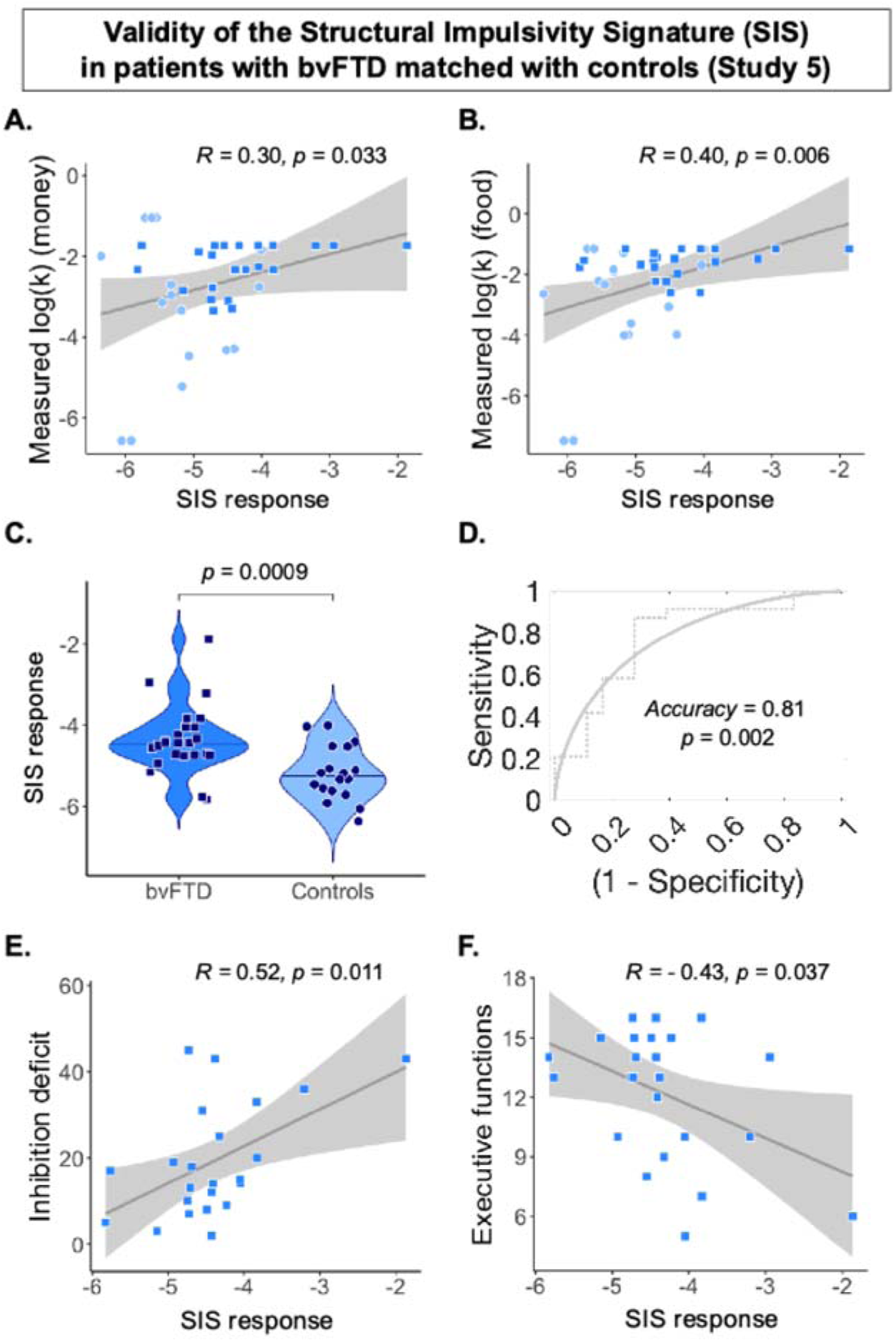
Predictive validity of the structural brain pattern in Study 5. **(A)** Correlation between SIS response and measured *log(k)* assessed with monetary rewards in Study 5 (*R*=0.30, *p*=0.07, 95%-*CI*= [-0.02, 0.57]). Patients are represented as squares in darker blue and controls as circles in lighter blue. **(B)** Correlation between SIS response and measured *log(k)* assessed with food rewards in Study 5 (*R*=0.45, *p*=0.01, 95%-*CI*= [0.1, 0.64]). Patients are represented as squares in darker blue and controls as circles in lighter blue. **(C)** As expected, SIS response was higher in bvFTD patients (*N*=24) than in controls (*N*=18) (*t*(40)=3.60, *p*=0.0009, Cohen’s *d*=1.09, 95%-*CI*=[0.41, 1.76]). **(D)** ROC curve showing the performance of SIS response in classification of bvFTD patients versus healthy controls (single interval test thresholded for optimal accuracy: accuracy=81 %, *p*= 0.002, *AUC*= 0.80, sensitivity = 87.5%, specificity = 72.2%). (**E)** Higher SIS responses were related to greater inhibition deficits (Hayling-error score) in bvFTD patients (*R*=0.52, *p*=0.01, 95%-*CI*= [0.14, 0.77]). **(F)** Higher SIS responses were related to more impaired executive functions (as measured with the FAB score) in bvFTD patients (*R*=-0.43, *p*=0.04, 95%-*CI*= [-0.71, -0.03]).

As expected, we found that SIS response was significantly higher in bvFTD patients than in controls (*t*(40)=3.60, *p*= 0.0009, Cohen’s *d*=1.09, 95%-*CI*= [0.41, 1.76], see Figure 4.C). Notably, SIS response significantly predicted whether a grey matter density map was from a bvFTD patient or from a control participant, with a classification accuracy of 81 % (*p*= 0.002, sensitivity = 87.5%, specificity = 72.2%, - see Figure 4.D).

Within the group of bvFTD patients, higher SIS response was associated with higher inhibition deficit (higher Hayling-error score; *R*=0.52, *p*=0.01, 95%-*CI*= [0.14, 0.77]) and higher executive troubles (lower FAB score; *R*=-0.43, *p*=0.04, 95%-*CI*= [-0.71, - 0.03]) (see Figure 4.E and 4.F). Importantly, we found that SIS responses remained significantly associated with impaired inhibition—as reflected by higher Hayling-error scores—within the bvFTD patient group, even after controlling for dementia severity (as measured by the Mattis Dementia Rating Scale; *B*=8.63, *p*=0.02, 95%-*CI*= [1.51, 15.7]) and disease duration (*B*=9.41, *p*=0.005, 95%-*CI*= [3.14, 15.7]). These results indicate that the predictive value of the brain marker for inhibitory dysfunction is not merely a reflection of overall disease severity or progression. Taken together, these findings show that the SIS significantly and accurately classified bvFTD patients from matched controls, and that it tracked the severity of key symptoms in these patients.

### Spatial distribution of weights in the SIS

#### Thresholded pattern of the structural brain signature

Bootstrapping results revealed the positive and negative weights that most strongly contributed to GMD-based prediction of individual differences in impulsivity (measured by delay discounting in Study 1). At a threshold of *q* = 0.05 (FDR-corrected), we identified two clusters where greater grey matter density was positively associated with individual differences in discounting—that is, higher grey matter density was linked to greater impulsivity. These clusters were located in the left lateral parietal cortex (supramarginal gyrus) and the left lateral occipital cortex (superior division). For exploratory purposes, using a threshold of *p* = 0.001 (uncorrected), we identified additional clusters with positive weights—primarily in regions associated with the valuation system(Lempert et al., 2019), including the right orbitofrontal cortex (OFC), ventromedial prefrontal cortex (vmPFC), and right ventral striatum.

At a threshold of *q* = 0.05 (FDR-corrected), we identified one cluster in the posterior cingulate cortex (PCC) and adjacent lingual gyrus—including the retrosplenial cortex—where grey matter density was negatively associated with discounting differences; that is, lower grey matter density in this region was linked to higher impulsivity. At a more liberal, exploratory threshold of *p* = 0.001 (uncorrected), additional regions showing negative weight contributions included the left hippocampus, right anterior insula (AI), dorsal anterior cingulate cortex (ACC), and bilateral amygdala. For visualization purposes, the bootstrapped weight map is presented in Figure 5A using a more inclusive threshold (*p* = 0.05 uncorrected; see also Supplementary Table 2).

**Figure 5.**
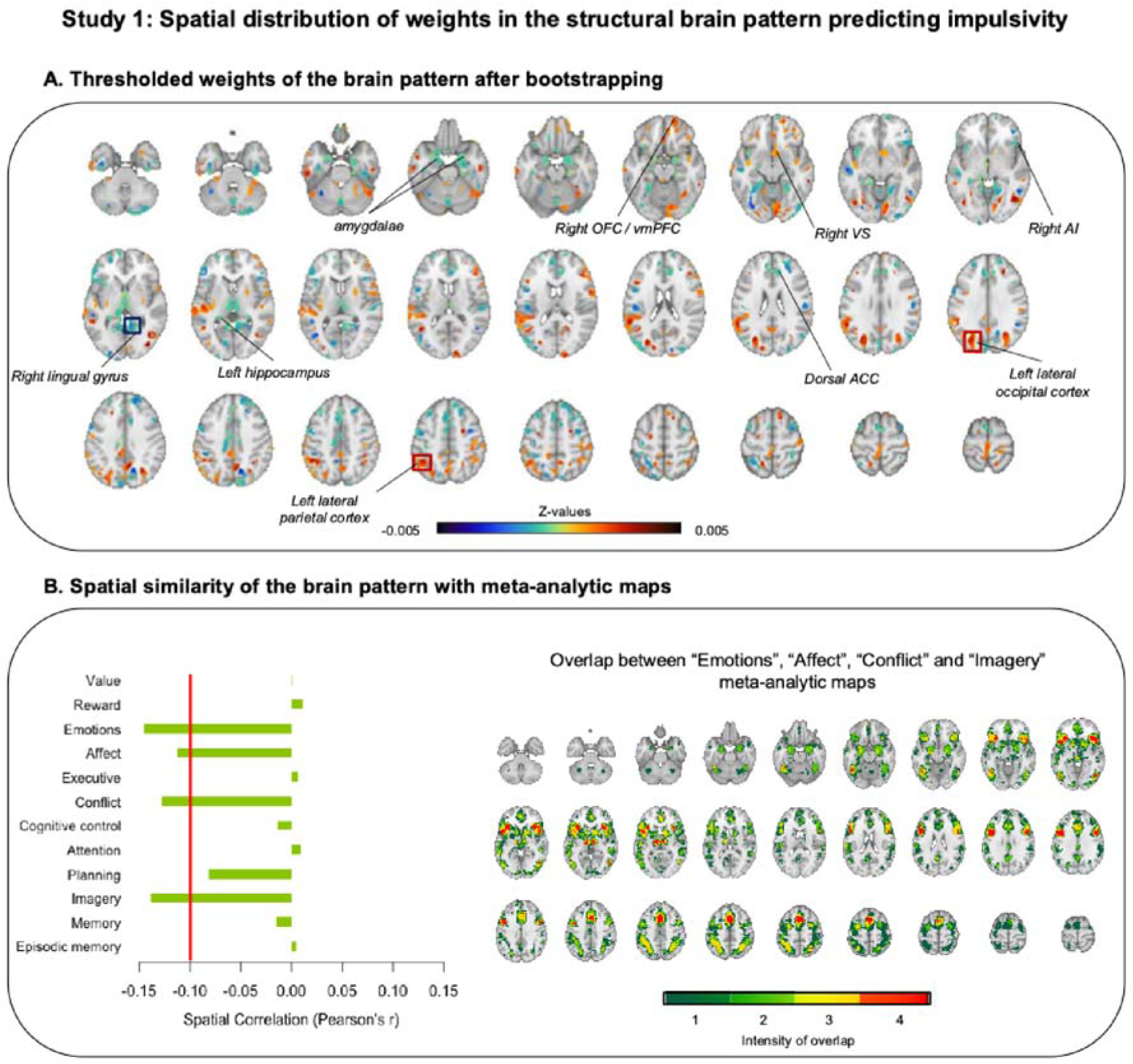
Spatial organization of the structural brain pattern developed in Study. **1. (A)** Whole-brain weight map thresholded at *p*=0.05 (uncorrected for multiple comparisons across the brain) resulting from a bootstrapping procedure (5,000 samples); negative weights (contributing to lower impulsivity with higher grey matter density) are shown in blue. Positive weights (contributing to higher discounting with higher grey matter density) are shown in orange. The three framed clusters correspond to the three clusters in which peaks are significant at *q*=0.05 FDR-corrected. Regions indicated in italics are some of the main regions significant at *p*=0.001, uncorrected (OFC: orbitofrontal cortex; vmPFC: ventromedial prefrontal cortex; VS: ventral striatum; AI: anterior insula; ACC: anterior cingulate cortex). **(B)** On the left, spatial correlations of the unthresholded impulsivity brain pattern with thresholded meta-analytic uniformity maps from Neurosynth (http://www.neurosynth.org). As in(Koban et al., 2023), we selected meta-analytic maps corresponding to three types of functions assumed to be involved in delay discounting: 1/ valuation and emotion processing; 2/ executive control; 3/ memory and prospection. Spatial correlations are descriptive and indicate the extent of spatial similarities between the structural brain pattern and the functional networks of interest(Yarkoni et al., 2011). Highest correlations (or similarities) were observed with the “Emotions”, “Affect”, “Conflict”, and “Imagery” meta-analytic maps, and were all negative, meaning that higher grey matter density in these functional regions is associated with lower impulsivity. On the right, we show the spatial distribution and overlap between the four meta-analytic maps found to be the most negatively correlated with the structural brain pattern (from 1, corresponding to non-overlapping regions from only one map, to 4, corresponding to regions of overlap between the 4 maps).

### Similarity of the structural brain signature to meta-analytic maps

When comparing the predictive map of impulsivity with meta-analytic uniformity maps (Yarkoni et al., 2011) (representing functional networks that have been previously hypothesized (Peters and Büchel, 2011) to contribute to impulsivity), we observed that the highest similarities (spatial correlation *r*’s > 0.1 in absolute value) were with the “Emotions”, “Affect”, “Conflict” and “Imagery” meta-analytic maps (Figure 5B, left panel). These spatial correlations were all negative, indicating that greater grey matter density in areas related to emotions, affect, conflict processing, and imagery contributes to predicting lower impulsivity (or conversely, lower grey matter density in these areas predicts higher impulsivity). The “Emotions”, “Affect”, “Conflict” and “Imagery” meta-analytic maps correspond to overlapping functional networks (see Figure 5B, right part). Among the most overlapping regions between these four networks (in red), the AI and dorsal ACC, corresponding to robust negative weights in the brain pattern, are known to be major hubs of the salience network (Menon, 2015).

### Spatial distribution of brain regions contributing to higher SIS responses in bvFTD

For each bvFTD patient and each control participant, we computed an ‘importance map’ as the unsummed matrix dot product between the predictive structural weight map and the individual grey matter density map. Since higher resulting dot product contributes to higher SIS response, the importance map shows which brain regions contributed to increase (or decrease) SIS response in each individual. We performed a t-test contrasting bvFTD patients and controls (bvFTD > controls) on the resulting importance maps, with a family-wise error (FWE) correction applied to p-values to correct for multiple comparisons across the brain (see Figure 6C). This contrast shows the regions in which structural atrophy contributed positively to higher SIS response in bvFTD than in controls (regions in red). These included the OFC, anterior insula, dorsal ACC, striatum, thalamus, amygdala, hippocampus, and middle temporal regions. These regions corresponded to areas combining the presence of negative weights in the predictive brain pattern (i.e., voxels for which higher GMD predicts lower impulsivity, shown in Figure 6B) and the presence of significant grey matter atrophy due to bvFTD pathology (see atrophy pattern in Figure 6A). Thus, the contrast shown in Figure 6C also maps the regions in which the SIS is the most similar to bvFTD atrophy pattern.

**Figure 6.**
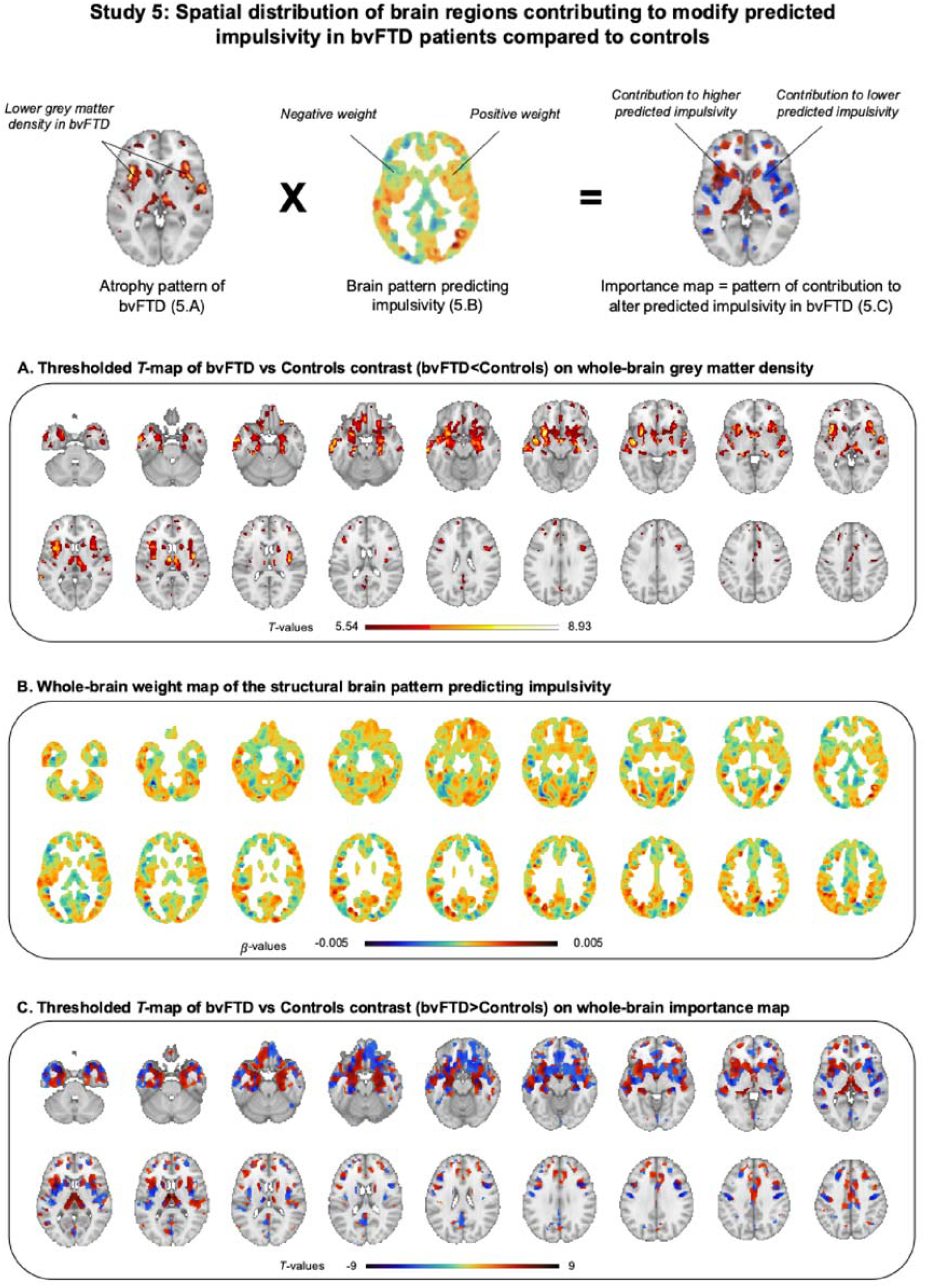
Spatial distribution of regions contributing to higher SIS response in bvFTD in Study 5. We computed an importance map as the unsummed matrix dot product between the Structural Impulsivity Signature (SIS) (developed in Study 1) and the individual grey matter density map of each Study 3 participant. Since higher resulting dot product contributes to higher SIS response, the importance map shows how brain regions contribute to increased (or decreased) SIS response in each individual. We performed a t-test contrasting bvFTD patients and controls (bvFTD > controls) on the resulting importance maps, to show the regions in which the contribution to higher SIS response was significantly higher in bvFTD than in controls. Within regions showing atrophy in bvFTD (see 6.A), those corresponding to negative (/positive) weights in the whole-brain predictive pattern (see 6.B) contributed to increase (/decrease) SIS response in bvFTD (see 6.C). **(A)** VBM–derived grey matter atrophy map of bvFTD patients contrasted with matched controls (bvFTD<Controls), FWE-corrected and thresholded at *p* < 0.05. **(B)** Unthresholded whole-brain weight map of the structural brain pattern developed in Study 1 and used in Study 2 to predict impulsivity in bvFTD patients (*N*=24) and matched controls (*N*=18). Negative weights (contributing to lower impulsivity with higher grey matter density) are in blue and positive weights (contributing to higher impulsivity with higher grey matter density) are in orange. **(C)** Contrast between bvFTD patients and controls (bvFTD>Controls)) on the importance map, FWE-corrected and thresholded at *p* < 0.05; this map shows regions in which atrophy contributed to increased SIS response in bvFTD patients (compared to controls) in red and regions contributing to decrease SIS response in bvFTD patients (compared to controls) in blue, the balance being in favor of a global increase in SIS response in bvFTD patients.

## Discussion

Impulsive and maladaptive behaviors are transversal features of many neuropsychiatric disorders and especially prominent in behavioral-variant frontotemporal dementia (bvFTD). Yet, their relationship with individual brain structure is still debated. Here, we used a machine learning technique to develop a brain signature predicting individual differences in impulsivity based on whole-brain grey matter density patterns in a first sample of 117 healthy adults (Study 1). We further tested the generalizability of the obtained brain signature in four additional independent studies in healthy and clinical participants (Studies 2, 3, 4 and 5; total *N* = 626) and its clinical value in bvFTD patients (Study 5). We found that: 1) individual differences of whole-brain grey matter density reliably predicted individual differences in discounting rates and trait impulsivity in Study 1; 2) the resulting Structural Impulsivity Signature (SIS) predicted trait impulsivity in Studies 2, 3 and 4; 3) the SIS separated bvFTD patients from controls with 81% accuracy and significantly predicted differences in two disease-related impulsivity symptoms (i.e., inhibition deficit and executive deficits) among bvFTD patients.

The SIS predicted four different types of measures of impulsivity in different population samples: 1) a task-based behavioral measure (i.e., delay discounting) in healthy adults (Study 1) and across bvFTD patients and controls (Study 5); 2) a questionnaire-based personality measure of urgency from UPPS scale in healthy adults (Studies 1 and 2); 3) a questionnaire-based personality measure of impulsiveness from BIS scale across healthy and neuropsychiatric conditions (Studies 3 and 4); 4) clinical measures of core impulsivity symptoms of bvFTD from the Hayling test and Frontal Assessment Battery (in Study 5). This result is in agreement with the idea of the existence of one coherent latent concept of impulsivity which can be revealed by different types of measures (Chamberlain et al., 2018; Huang et al., 2024) and which constitutes a continuum across healthy and neurological conditions. All the measures predicted by the SIS are related to a concept of impulsivity defined as a general tendency to act in a rush (especially in response to an emotional state), selecting the most prepotent immediate answers to a given situation and disregarding potential consequences (Webster and Jackson, 1997).

The SIS provides insight into the neurobiological substrates of impulsivity. Regions previously associated with delay discounting and impulsivity (such as the OFC and vmPFC) (Ballard and Knutson, 2009; Cooper et al., 2013; Lebreton et al., 2013; Li et al., 2013; Hare et al., 2014; van den Bos et al., 2014; Pehlivanova et al., 2018; Koban et al., 2023; Bergström et al., 2024) did not survive conservative corrected thresholds after bootstrapping but they were among the major positive contributors to individual differences in impulsivity (relatively to other brain regions which did not survive the uncorrected thresholding of *p*=0.001). In addition, our whole brain approach uncovered the role of other regions (e.g., lateral parietal cortex and occipital cortex) not typically linked to impulsivity but consistent with previous findings regarding the neuroanatomical correlates of trait impulsivity measured by the BIS scale(Ide et al., 2017). Among the strongest negative contributors (in which lower grey matter density contributes to higher individual impulsivity), we found clusters corresponding to hub regions of the salience network (anterior insulae, dorsal ACC, amygdalae). These regions are primarily targeted by the neurodegenerative process in bvFTD patients (Seeley et al., 2009), thus explaining the higher impulsivity in these patients compared to controls. These regions are associated with the processing of emotionally significant internal and external stimuli (Seeley et al., 2012; Menon, 2015; Seeley, 2019) and awareness of present and future affective states (Craig, 2009); they are involved in switching between large-scale networks to facilitate access to attention and working memory resources in the presence of a salient event(Menon and Uddin, 2010). These areas are further known to be involved in cognitive conflict processing (Botvinick et al., 2001; Van Veen et al., 2001) and choice conflicts between different intertemporal options (Hoffman et al., 2008).

Because the SIS predicts core symptoms of bvFTD, it has the potential to contribute to the early diagnosis of bvFTD. Brain signatures can contribute to the diagnosis of conditions involving distributed and subtle brain lesions that are sometimes difficult to detect by mere visual inspection of MRI scans, especially at early stages of the disease. The SIS may contribute to the diagnosis of bvFTD by complementing other brain models able to detect bvFTD. A few previous studies successfully trained structural MRI classifiers for the specific purpose of distinguishing FTD patients from controls (e.g.,(Davatzikos et al., 2008; Gonzalez-Gomez et al., 2022; Moguilner et al., 2023)). These bvFTD classifiers have shown good accuracy to detect patients with clear structural brain damage but their ability to distinguish individuals at risk of developing FTD due to genetic mutations is likely to be limited to the period just before symptom onset (Feis et al., 2018, 2019). Under the hypothesis of a continuum of marked impulsivity in presymptomatic individuals and patients (Godefroy et al., 2023), the SIS might serve the early prediction and monitoring of bvFTD before symptom onset. Impulsive behaviors may be present in an attenuated form long before clinical diagnosis and hard to detect with traditional clinical methods. A neuromarker predicting impulsivity may be sensitive to specific brain modifications that appear early in individuals predisposed to FTD (possibly as neurodevelopmental lesions (Lee et al., 2017)) and would thus allow to enhance the monitoring of clinical signs of subtle behavioral changes. BvFTD patients in Study 5 were at a rather early stage of disease given their average MMSE score and disease duration. This suggests that the SIS is sensitive to early bvFTD. Future tests in genetic FTD populations are needed to evaluate the potential of the SIS to detect FTD even before symptom onset.

As it predicts nearly 30% of the variance of inhibition deficit among bvFTD patients, the SIS may be sensitive to lesions in a structural network underlying the core bvFTD symptom of disinhibition. In addition to its potential contribution to the early detection of presymptomatic individuals, this brain signature may thus provide insight into the neuropsychological profiles of patients. The SIS could become a useful tool to disentangle the phenotypic heterogeneity within bvFTD population (O’Connor et al., 2017). The characterization of different clinical and behavioral profiles within the bvFTD spectrum could help to better understand the pathology, and to better adapt treatments according to patients’ specific needs. The SIS predictions are also sensitive to individual differences in impulsivity among patients with different psychiatric conditions; this further supports the idea that the SIS could be a first step towards precision medicine neuromarkers to better define a patient’s profile. Additionally, future studies could test whether the SIS can distinguish bvFTD from other neurodegenerative or neuropsychiatric conditions which do not display impulsivity as core symptoms (e.g., Alzheimer’s disease). Using neuromarkers such as the SIS in cases of diagnostic uncertainty could help in the choice of treatment (Sheikh-Bahaei et al., 2017).

### Limitations

Our results point at challenges in generalizing brain signatures to other independent samples, which is crucial to establish boundary conditions in this rapidly growing field. We were successful at predicting delay discounting from whole-brain grey matter in a first sample of healthy adults showing a positive correlation between the discounting rate and urgency. In a second independent sample of healthy adults with no significant correlation between the discounting rate and urgency, we could not replicate the significant association with measured discounting rates. In the sample of Study 2, the theoretical relationship between measures of urgency and delay discounting is not confirmed, which implies difficulties to validate SIS responses with both of these measures. The discounting rate is influenced by the latent trait impulsivity but also depends on many other factors (Odum and Baumann, 2010) such as cultural context (Kim et al., 2012), social context (Schwenke et al., 2022) as well as participants’ cognitive abilities (Hirsh et al., 2008) (in Study 2, the correlation of delay discounting with IQ is *R*=-0.22; *p* = 0.004). However, we still found evidence of the conceptual validity of SIS responses in Study 2 through their significant link with reliable questionnaire measures (Enkavi et al., 2019) of trait urgency. Our findings suggest that the SIS most reliably predicts the measures of impulsivity traits by self-report questionnaires, which might be the most stable ones (as compared to other task-based behavioral measures) (Enkavi et al., 2019).

Although multivariate brain signatures can be replicable with moderate sample sizes (Spisak et al., 2022), future studies aimed at developing brain signatures of impulsivity could also benefit from using even larger and more diverse samples (Marek et al., 2022). We note that our results suggest a relatively small contribution of interindividual variability in brain structure to interindividual variability in impulsivity among healthy adults (*R^2^*=12.3% explained variance as a maximum in Study 1). These effect sizes are however in line with those reported for most brain signatures of behavioral individual differences using structural features (Genon et al., 2022). Of note, variability in genotype accounts for a similar part of the variance of impulsivity (Sanchez-Roige et al., 2018). The magnitude of associations between brain structure and behaviors may be limited in the general population but these associations might be more salient within populations with a marked variability of both brain and behavior such as patients with neurodegenerative conditions (about 30% explained variance among bvFTD patients).

## Conclusions

Our results reveal a distributed pattern of gray matter density to predict individual differences in impulsivity in healthy and neurological populations. This novel brain signature, the Structural Impulsivity Signature (SIS), further distinguished patients with bvFTD—a neurodegenerative disorder characterized by high impulsivity and maladaptive decision-making—from matched controls. While a few previous studies have developed functional brain signatures of delay discounting or impulsivity (Koban et al., 2023; Bergström et al., 2024), these remain dependent on functional (task-related) MRI data and have not yet been tested in neurological conditions. By identifying a structural network associated with individual differences in impulsivity across healthy and clinical populations, our results provide insight into the neurobiological bases of trait impulsivity and suggest a continuum of brain structure-impulsivity relationship. Our results also highlight the potential clinical utility of the SIS for diagnosing bvFTD, particularly in aiding the stratification of this heterogeneous condition. MRI could be instrumental to confirm an FTD diagnosis (Sheikh-Bahaei et al., 2017) and the SIS only requires a preprocessed T1-weighted scan to reach a prediction. Future studies are needed to fully test the clinical potential of the SIS in bvFTD and other neurological or neuropsychiatric conditions. Moreover, the SIS has the potential to be a key tool for uncovering how lasting structural changes in the brain link adverse experiences and environmental influences to increased impulsive decision-making.

## Acknowledgements

We thank all the participants and organizers of the five studies mentioned in this paper as well as all the students, in particular Daniela Schelski (for Study 1), who contributed to data collection.

