## Supplementary for "A structural MRI marker predicts individual differences in impulsivity and classifies patients with behavioral-variant frontotemporal dementia from matched controls"

**Additional Methods:** **MRI data preprocessing**

For Studies 1, 3, 4 and 5, we used the SPM module “Segment” for segmentation, bias correction and rigid alignment of T1 images. These images were then used as input into the DARTEL SPM module to create a customized DARTEL template and individual ‘flow fields’ for each subject. DARTEL determines the nonlinear deformations for warping all grey and white matter images so that they match each other. Finally, the SPM module “Normalise to MNI space” generated spatially normalized grey matter images using the deformations estimated in the previous step and images were spatially smoothed with a 6 mm Gaussian FWHM kernel.

For Study 2, T1 images were previously preprocessed for VBM analyses using the default preprocessing pipeline of the Computational Anatomy Toolbox (CAT12) for SPM12. T1-weighted images underwent spatial adaptive non-local means (SANLM) denoising filter, were bias corrected, and affine-registered, followed by standard SPM unified tissue segmentation into grey matter, white matter, and cerebral spinal fluid. The grey matter volume images were spatially registered to a common template using Geodesic Shooting, resampled to 1.5 mm^3^, and spatially smoothed with an 8 mm Gaussian FWHM kernel.

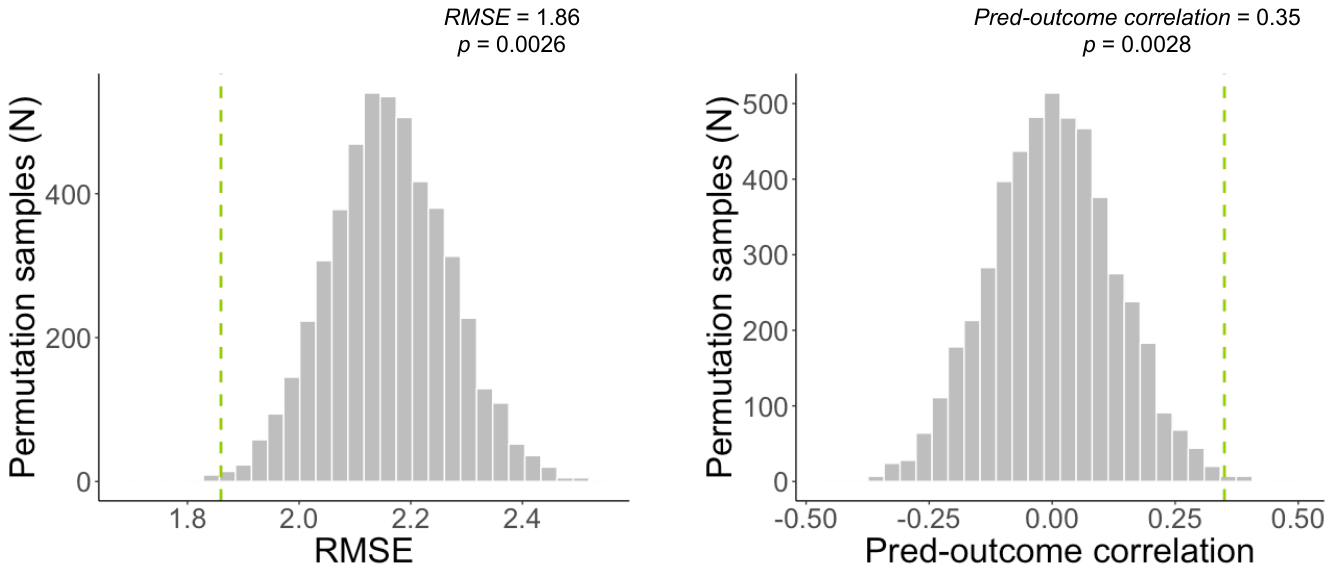

**Supplementary figure 1. Results of the permutation test (Study 1).** Delay discounting values were randomly permuted, and predictions were generated from 5000 randomly permuted samples to provide null distributions for standard metrics assessing the accuracy of prediction. The mean squared error (MSE) and mean absolute error (mean abs error) are shown in Figure 2. Here, we show results for complementary metrics: root mean squared error (RMSE) and prediction-outcome correlation. Histograms of null distribution are shown in gray bars, and the observed metrics are indicated by the green dotted lines.

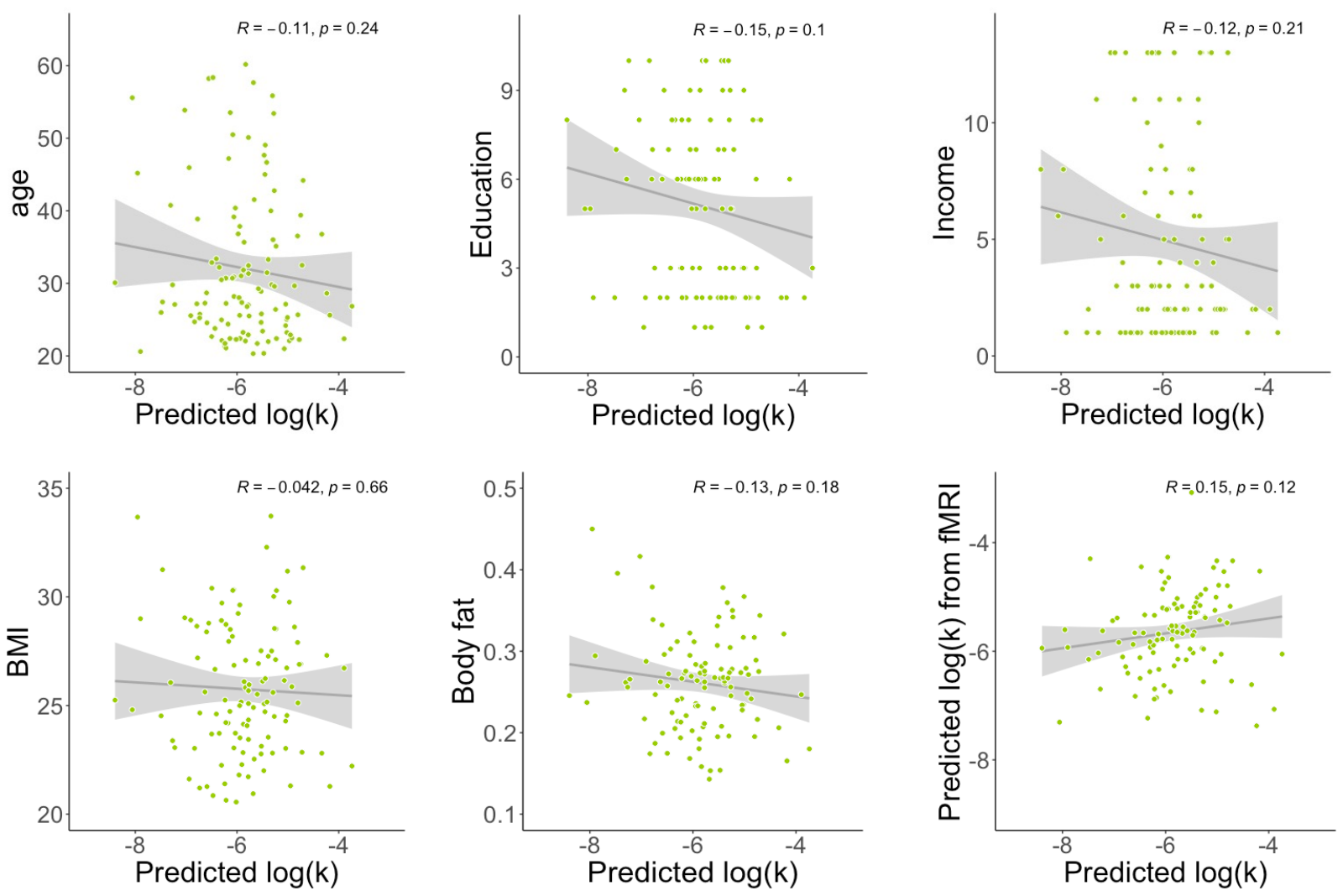

**Supplementary figure 2. Correlations between impulsivity predicted by the structural brain pattern and other individual characteristics in Study 1.** The selected characteristics of interest are demographic variables and BMI / body fat which have previously been related to delay discounting. Moreover, we tested the correlation between individual brain responses of the fMRI-based brain marker of log(k) and individual brain responses of the structural MRI-based brain marker of log(k) (see bottom right-hand corner).

**Supplementary Table 1**. **Demographical and main neuropsychological measures of bvFTD patients and controls.** Data are given as Mean (SD). BvFTD patients: N= 24 / Controls: N= 18; FAB: Frontal Assessment Battery; Hayling – error: objective measure of inhibition deficit from the Hayling Sentence Completion Test; MMSE: Mini-Mental State Examination; DRS: Mattis Dementia Rating Scale; SAS: Starkstein Apathy Scale; HADS: Hospital Anxiety and Depression Scale.

|  | bvFTD | Controls | bvFTD vs controls |
| --- | --- | --- | --- |
| % Women | 33.3% | 55.6% | $\chi^{2}$ *=* 1.3 ; *p* = 0.26 |
| Age | 66.6 (8.3) | 62.6 (7.2) | *t*(40) *=* 1.6 ; *p* = 0.11 |
| Education level | 6.3 (1.9) | 7.2 (1.0) | *t*(40) = -2.0 ; *p* = 0.05 |
| Hayling - error | 19.2(13.2) | 3.1(2.6) | *t*(40) = 5.7 ; *p* < .001 |
| FAB (/18) | 12.2 (3.3) | 17.3 (0.8) | *t*(40) = -7.3 ; *p* < .001 |
| MMSE (/30) | 23.8 (2.7) | 29.4 (0.8) | *t*(40) = -9.7 ; *p* < .001 |
| DRS (/144) | 118.5 (9.1) | 142.2 (1.3) | *t*(40) = -12.6 ; *p* < .001 |
| SAS (/42) | 16 (4.7) | 5.7 (3.1) | *t*(40) *=* 8.5 ; *p* < .001 |
| HADS-Anxiety (/21) | 7.7 (4.6) | 4.2 (2.4) | *t*(40) *=* 3.2 ; *p* = 0.003 |
| HADS-Depression (/21) | 1.2 (1.0) | 5.8 (3.2) | *t*(40) *=* 6.5 ; *p* < .001 |

**Supplementary Table 2. Significant positive and negative weights contributing to the structural brain marker predicting delay discounting in Study 1.** Only FDR-corrected clusters (q < 0.05 across the whole brain), with at least three contingent voxels are listed. X, Y, Z are the MNI coordinates of the local maxima. In the column ‘Atlas label’, regions are labeled based on Harvard-Oxford structural atlas. Most important significant clusters (in terms of size) are indicated in italics.

***Positive weights***

| **Atlas label** | **X** | **Y** | **Z** | **Volume (voxels)** | **max(Z)** |
| --- | --- | --- | --- | --- | --- |
| *Right lingual gyrus* | 14 | -51 | -2 | 96 | -0.0023203 |
| Right precuneous cortex | 16 | -66 | 38 | 8 | -0.0034364 |
| Right Frontal pole | 28 | 46 | 38 | 3 | -0.0016211 |

***Negative weights***

| **Atlas label** | **X** | **Y** | **Z** | **Volume (voxels)** | **max(Z)** |
| --- | --- | --- | --- | --- | --- |
| *Left lateral occipital cortex* | -30 | -74 | 28 | 82 | 0.0032081 |
| *Left supramarginal gyrus* | -46 | -48 | 45 | 36 | 0.0032853 |
| Left supramarginal gyrus | -58 | -44 | 20 | 11 | 0.0027848 |
| Right lingual gyrus | 12 | -78 | -9 | 7 | 0.0022673 |
| Right precuneous cortex | 10 | -56 | 36 | 3 | 0.0031411 |
| Precentral gyrus | 0 | -22 | 69 | 3 | 0.0014893 |
